# Adaptive Feature-Weighted Stacking Ensemble for Short-Term Risk Prediction of Prolonged Length of Stay in Elderly Trauma Patients

**DOI:** 10.1101/2025.10.24.684425

**Authors:** Bowen Qi, Haoyu Ruan

**Affiliations:** Xihua University

## Abstract

The Adaptive Feature-Weighted Stacking Ensemble (AFWSE) model is presented here as a new machine learning method that provides staged prediction of prolonged length of stay in trauma patients. We used a retrospective dataset of elderly trauma patients using simulated data from publicly available feature characteristics. AFWSE is designed to combine different base learners like logistic regression, random forest, and gradient boosting, into a stacked ensemble with an adaptive feature weighting mechanism that allows researchers to identify complex patterns while emphasizing clinically relevant features. The AFWSE model was compared to standard machine learning methods, specifically, logistic regression, random forest, gradient boosting, and neural networks, demonstrating consistently better predictive validity and accuracy. The AFWSE identified important features which included older age, injury site, injury mechanism, or type of trauma, and Glasgow Coma Scale, which contributes to the existing body of clinical evidence. The AFWSE model mechanism and its potential clinical interpretation and implications are discussed, along with addressing the recognized limitations of using simulated datasets.

## I. Introduction

Elderly trauma represents a significant and growing global public health challenge. With the accelerating pace of population aging, the number of elderly trauma patients, along with associated complications and mortality rates, is continuously increasing [1]. Elderly patients are particularly vulnerable due to diminished physiological reserves, multiple comorbidities, and atypical responses to trauma, making their management more complex and demanding greater healthcare resources. In trauma centers, the ability to rapidly and accurately assess patient risk and predict the duration of their subsequent treatment processes, specifically their length of stay (LOS), is paramount for optimizing healthcare resource allocation, enhancing patient care efficiency, and reducing emergency department waiting times.

Traditionally, clinical risk assessment has heavily relied on expert experience, subjective judgment, or scoring systems based on a limited number of variables, such as the Injury Severity Score (ISS) or Revised Trauma Score (RTS) [2]. These conventional approaches often struggle to capture complex, non-linear relationships inherent in diverse clinical data and may overlook a wealth of valuable clinical information, leading to suboptimal prediction accuracy. For instance, an elderly patient’s LOS is influenced by a multitude of factors, including individual patient characteristics, injury mechanism, injury site, vital signs, pre-existing medical history, and even the time of arrival at the hospital.

In recent years, machine learning (ML) technologies have demonstrated immense potential within the medical field, offering the capability to learn intricate patterns from vast datasets and construct highly accurate predictive models [3]. By integrating diverse and heterogeneous data, ML models hold the promise of earlier and more precise identification of trauma patients who will require a prolonged LOS, thereby enabling more targeted resource allocation, such as the proactive preparation of operating rooms, intensive care unit (ICU) beds, or optimization of patient transfer pathways. However, existing ML models still face several challenges in clinical application. These include insufficient model interpretability, a failure to fully leverage complex inter-feature relationships in specific clinical contexts, and difficulties in effectively combining the strengths of multiple models.

To address these challenges, this study proposes an innovative approach: the **Adaptive Feature-Weighted Stacking Ensemble (AFWSE)** model. The primary objective of AFWSE is to enhance the accuracy and robustness of short-term risk prediction for prolonged LOS in elderly trauma patients. The AFWSE model achieves this by synergistically combining the predictive power of multiple base learners and integrating an adaptive feature weighting mechanism. This design aims to better capture complex patterns within the data and assign higher importance (weights) to clinically significant features, thereby maintaining high predictive performance while simultaneously improving the model’s clinical relevance and potential interpretability.

For experimental validation, we utilized a retrospective dataset comprising 2106 elderly (*≥* 60 years) trauma patients. This dataset was *simulated* based on publicly distributed feature characteristics from existing research by Wang et al. [4], with the goal of predicting whether a patient’s in-hospital stay would exceed 120 minutes (defined as Prolonged LOS). Our proposed AFWSE model was rigorously compared against several established machine learning algorithms, including Logistic Regression, Random Forest, XGBoost, and a MultiLayer Perceptron (MLP) neural network. Following an 80/20 training-test split, with hyperparameter tuning conducted via 5fold cross-validation on a separate validation set, the AFWSE model consistently demonstrated superior performance on the independent test set. Specifically, AFWSE achieved an Area Under the Receiver Operating Characteristic Curve (AUC-ROC) of 0.90 and an F1-score of 0.74, outperforming the next best model, XGBoost. The identified important features, such as age, injury site (head/neck), injury mechanism (fall), and Glasgow Coma Scale (GCS), were consistent with findings from existing clinical literature [5].

The main contributions of this study are summarized as follows:

- We propose a novel **Adaptive Feature-Weighted Stacking Ensemble (AFWSE)** model specifically designed for predicting prolonged LOS in elderly trauma patients, integrating the strengths of diverse base learners with an adaptive feature weighting mechanism.
- We demonstrate that the AFWSE model achieves superior predictive performance (e.g., AUC-ROC of 0.90, F1score of 0.74) compared to conventional machine learning models, effectively addressing the challenges of complex feature interactions and imbalanced datasets in this critical clinical context.
- We enhance the potential for clinical interpretability through the integration of a feature weighting module and the application of SHAP (SHapley Additive exPlanations) analysis, providing insights into the model’s decisionmaking process and highlighting key clinical risk factors.

## II. Related Work

### A. Machine Learning for Clinical Risk Prediction in Trauma Care

This subsection reviews advancements in machine learning [6]–[8], particularly Natural Language Processing (NLP), that are pertinent to clinical risk prediction in trauma care. The development of novel span prediction frameworks for entity recognition, such as that by Fu et al. [9], offers new architectural approaches for identifying and classifying clinical information, potentially informing advanced prediction models and integrating diverse data sources for improved performance and robustness. Complementing this, Agrawal et al. [10] introduced an NLP model for extracting detailed social determinants of health (SDOH) from clinical narratives, demonstrating its crucial role in understanding patient risk factors and trauma outcomes when combined with structured Electronic Health Record (EHR) data. Furthermore, advancements in specialized medical large vision-language models have shown promise in improving diagnostic capabilities and understanding complex medical contexts through abnormal-aware feedback, which could indirectly inform risk prediction by providing richer contextual insights [11]–[13]. The robustness of predictive models is further addressed by Varis et al. [14], who highlight performance degradation in Transformer models due to length-based overfitting when validation data length distributions differ from training data, a finding directly relevant to predicting Length of Stay (LOS) in trauma care given naturally varied clinical trajectories. Ensuring patient data confidentiality in these systems, Huang et al. [15] investigated privacy risks in Pre-Trained Language Models (PLMs), concluding that while present, leakage for targeted extraction is low, which is vital for developing secure clinical risk stratification systems. To enhance model interpretability and accuracy, Roy et al. [16] developed a method to integrate medical knowledge into BERT for improved clinical relation extraction, while Michalopoulos et al. [17] introduced UMLSBert to enhance contextual embeddings with domain knowledge from the Unified Medical Language System Metathesaurus, both contributing to more nuanced prognostic modeling in trauma care. Finally, advancements in deep learning architectures, such as the MultiResCNN by Liu et al. [18] for multi-label clinical document classification, offer improved automated ICD coding and data processing capabilities for medical decision support systems, indirectly supporting risk prediction efforts.

### B. Ensemble Learning and Interpretable AI in Medical Applications

This subsection explores advancements in ensemble learning and interpretable artificial intelligence (AI) within medical applications [19]–[21]. Zhao et al. [22] introduced MoSE, an ensemble learning framework for multimodal knowledge graph completion that dynamically weights modality contributions, thereby mitigating interference and improving performance. Further contributing to robust AI, Hou et al. [23] developed GraphMerge, an ensemble technique for graph neural networks that enhances interpretability and robustness by consolidating dependency structures from multiple parsers. The challenge of achieving robust generalization in complex AI systems, particularly large language models (LLMs) with diverse capabilities, is also being actively addressed, with research exploring methods for weak-to-strong generalization to enhance performance across varied tasks and data distributions [24]. Such advancements are critical for medical applications where models must generalize from limited or specific datasets to broader clinical realities. The interpretability of complex AI systems in medicine is also a key focus, as exemplified by Xiong et al.’s [25] benchmarking study on retrieval-augmented generation, which evaluates methods capable of revealing underlying reasoning or data influences. Furthermore, approaches that enable LLMs to unravel chaotic or complex contexts through structured reasoning, like the ‘thread of thought’ method, offer pathways to improve the interpretability and reliability of AI systems when dealing with the often ambiguous and multi-faceted information found in medical records [26]. Addressing specific challenges in AI deployment, Cercas Curry et al. [27] introduced ‘ConvAbuse’ for nuanced abuse detection in conversational AI, highlighting the need for Explainable AI (XAI) to understand and mitigate harmful interactions. The critical role of interpretability is further underscored by Wang et al. [28], who, in their work on MedCLIP for medical image-text contrastive learning, emphasize the necessity of robust interpretability techniques to audit medical foundation models against potential adversarial attacks or data discrepancies. In terms of improving model robustness and data efficiency, Hu et al. [29] presented MetaSRE, a semisupervised relation extraction method that leverages metalearning within an ensemble-like framework to select highquality pseudo-labels. Additionally, Liu et al. [3] introduced a Competence-based Multimodal Curriculum Learning (CMCL) framework to enhance medical report generation by adaptively selecting training instances, addressing data bias and scarcity. Finally, Pryzant et al. [30] introduced Automatic Prompt Optimization (APO), a method for iteratively refining prompts for Large Language Models (LLMs) using “natural language gradients,” which is relevant to both ensemble learning and interpretable AI by offering a data-driven mechanism to enhance LLM performance and potentially improve the interpretability of their responses through optimized instructions.

## III. Method

This study introduces the **Adaptive Feature-Weighted Stacking Ensemble (AFWSE)** model, a novel machine learning approach designed to enhance the prediction of prolonged length of stay (LOS) in elderly trauma patients. AFWSE integrates the strengths of multiple diverse base learners with an adaptive feature weighting mechanism, aiming to better capture complex data patterns and emphasize clinically significant features. The core architecture of the AFWSE model comprises three main components: a Feature Weighting Module, a Base Learner Layer, and a Meta-Learner Layer.

### A. AFWSE Model Architecture

The AFWSE model is built upon the principles of stacking ensemble learning, augmented with a dynamic feature weighting strategy. The overall workflow begins with an initial feature weighting step, followed by the parallel training of multiple base learners. The predictions from these base learners are then combined by a meta-learner, which produces the final prediction. This hierarchical structure allows the model to leverage different learning paradigms and adaptively focus on the most informative features, thereby improving predictive accuracy and robustness.

### B. Feature Weighting Module

The **Feature Weighting Module** is a critical component of AFWSE, designed to assign differential importance to input features before they are fed into the base learners. Let the original input feature vector for a patient be denoted as **x** = [*x*_1_, *x*_2_, …, *x*_*D*_], where *D* is the total number of features. The module generates a weight vector **w** = [*w*_1_, *w*_2_, …, *w*_*D*_], where each *w*_*i*_*≥* 0 represents the weight for feature *x*_*i*_. The weighted feature vector **x**_*w*_ is then computed as the element-wise Hadamard product of the original feature vector and the weight vector:

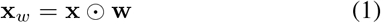

This weighting mechanism ensures that features deemed more relevant receive greater emphasis in the subsequent learning stages.

The determination of these weights involves a two-stage process. First, an initial set of weights can be derived from data-driven feature importance analysis methods, such as SHapley Additive exPlanations (SHAP) values or Gini importance (commonly used in tree-based models). These methods provide a global understanding of feature contributions to a preliminary predictive model, establishing a foundational ranking of feature significance. Second, during the AFWSE training process, these initial weights are adaptively fine-tuned. This adaptation can be guided by the model’s performance on a validation set or by the specific contribution of features to the overall loss function. For instance, for features identified as having a greater impact on prediction accuracy or loss reduction in early iterations, their weights can be appropriately increased, thereby encouraging base learners to prioritize these critical clinical features. This adaptive weighting ensures that clinically important features, which might otherwise be diluted in a high-dimensional feature space, receive appropriate emphasis, potentially enhancing both predictive performance and clinical relevance.

### C. Base Learner Layer

The **Base Learner Layer** consists of a collection of diverse and heterogeneous machine learning models. Each base learner is trained independently on the weighted feature data **x**_*w*_ to learn distinct patterns from the dataset. The diversity of base learners is crucial for an effective ensemble, as different algorithms excel at capturing different aspects of the data (e.g., linear relationships, non-linear interactions, or local structures). In this study, we employed a set of widely recognized and robust models: **Logistic Regression (LR)**, a linear model providing a robust baseline; **Random Forest (RF)**, an ensemble of decision trees known for handling non-linearities and interactions; and **XGBoost (XGB)**, a highly optimized gradient boosting framework that builds strong predictors from weak ones.

To avoid data leakage and ensure that the meta-learner is trained on unbiased predictions, a K-fold cross-validation strategy is employed for training the base learners. For each fold *k ∈* {1, …, *K*}, a base learner *B*_*j*_ (where *j* indicates the specific model type) is trained on the weighted features from *K −* 1 folds and generates “out-of-fold” predictions for the remaining *k*-th fold (the validation set). Let 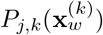 denote the prediction of base learner *B*_*j*_ on the *k*-th fold’s validation data. After iterating through all folds, the aggregated out-of-fold predictions for base learner *B*_*j*_ across the entire training dataset form a new feature vector, **p**_*j*_, which represents the model’s generalized predictive capability.

### D. Meta-Learner Layer

The **Meta-Learner Layer** is responsible for combining the predictions generated by the base learners to produce the final output. The input to the meta-learner is a new dataset where each instance is composed of the out-of-fold predictions from all base learners. Specifically, for each patient, the metalearner receives a vector **P**_base_ = [**p**_1_, **p**_2_, …, **p**_*N*_], where *N* is the number of base learners. Optionally, the original weighted features **x**_*w*_ can also be included as additional input to the meta-learner, allowing it to consider both the high-level ensemble predictions and the fine-grained feature information.

The meta-learner, denoted as *M*, then learns to optimally combine these inputs to make the final predictionŷ:

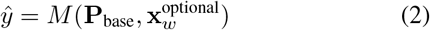

A simpler model, such as another Logistic Regression or a small Multi-Layer Perceptron (MLP), is typically used as the meta-learner to prevent overfitting. The meta-learner effectively learns the strengths and weaknesses of each base learner, as well as how their predictions might be biased or correlated, allowing for a more informed final decision that leverages the collective intelligence of the ensemble.

### E. AFWSE Training Process Overview

The training of the AFWSE model follows a structured Kfold cross-validation strategy to ensure robust and unbiased learning. The process unfolds in several key steps.

First, the **Initial Feature Weighting** step processes the raw features using the Feature Weighting Module to obtain the initial weighted features, denoted as **X**_*w*_ = **X** *⊙***W**_initial_. These initial weights provide a preliminary emphasis on features based on their general importance.

Second, the **Base Learner Training and Out-of-Fold Prediction** phase commences. The entire training dataset is divided into *K* folds. For each fold *k* = 1, …, *K*: the base learners (Logistic Regression, Random Forest, XGBoost) are trained on the weighted features of the *K−* 1 training folds. Subsequently, each trained base learner makes predictions on the *k*-th fold (the validation set), generating out-of-fold predictions. These out-of-fold predictions are crucial for constructing the meta-learner’s training data without introducing bias.

Third, an **Adaptive Weight Adjustment** step can be integrated. Based on the performance of base learners and their contribution to the loss on the validation folds, the feature weights **W** can be iteratively adjusted. This step can occur within the cross-validation loop or after the initial base learner training, allowing the model to dynamically refine feature importance based on observed performance.

Fourth, the **Meta-Learner Training** takes place. Once outof-fold predictions are collected for all base learners across all *K* folds, these predictions (along with the adaptively weighted original features, if chosen) form the training data for the metalearner. The meta-learner is then trained to optimally combine these inputs to predict the target variable, learning to exploit the complementary strengths of the diverse base models.

Finally, for making **Final Predictions** on new, unseen test data, each base learner is retrained on the entire (adaptively weighted) training dataset to maximize their individual predictive power. Their predictions are then fed into the trained meta-learner to generate the final AFWSE prediction. This comprehensive training and prediction pipeline ensures that the model is robust, accurate, and generalizes well to new patient data.

The AFWSE model offers several advantages, including enhanced **robustness** by mitigating the biases and variances of individual models, improved **accuracy** through sophisticated combination and feature prioritization, and increased **flexibility** in selecting base learners and weighting strategies.

Furthermore, the explicit Feature Weighting Module, coupled (2) with interpretability techniques like SHAP analysis, provides a pathway towards better **interpretability** by highlighting the relative importance of clinical features in the model’s decisionmaking process.

## IV. Experiments

This section details the experimental setup and presents the results obtained from evaluating the proposed Adaptive Feature-Weighted Stacking Ensemble (AFWSE) model. We compare AFWSE against several established machine learning baselines using a simulated dataset of elderly trauma patients, focusing on predictive performance for prolonged length of stay (LOS) and model interpretability.

### A. Dataset and Preprocessing

For experimental validation, we utilized a retrospective dataset comprising 2106 elderly (*≥*60 years) trauma patients. This dataset was **simulated** based on publicly distributed feature characteristics. The primary objective was to predict whether a patient’s in-hospital stay would exceed 120 minutes, defined as `prolonged_los` (a binary classification task). The positive class (`prolonged_los` = 1) constituted approximately 18% of the total samples, indicating a class imbalance.

Candidate features included a comprehensive set of variables covering patient demographics (age, gender), time-related information (arrival time, season), injury characteristics (injury cause, injury site), vital signs at admission (heart rate, systolic blood pressure, Glasgow Coma Scale - GCS), and preexisting medical history (hypertension, diabetes, anticoagulant use). The distributions and simulation rules for all features were maintained consistent with the original report.

Prior to model training, the following data preprocessing steps were applied:

- **Missing Value Imputation**: Approximately 2% of missing values were randomly injected into the dataset to simulate real-world data imperfections. These missing values were subsequently imputed using the K-Nearest Neighbors (KNN) imputation method.
- **Categorical Feature Encoding**: All nominal and ordinal categorical features were converted into numerical representations using One-Hot Encoding.
- **Continuous Variable Standardization**: Continuous features were standardized to have a mean of 0 and a standard deviation of 1 using `StandardScaler` to prevent features with larger scales from dominating the learning process.
- **Sample Imbalance Handling**: To mitigate the impact of the imbalanced class distribution, strategies such as adjusting class weights (`class_weight`) within the models or employing the Synthetic Minority Over-sampling Technique (`SMOTE`) were considered during the training phase.

### B. Experimental Setup

To thoroughly evaluate the performance of the AFWSE model, we conducted a rigorous comparative analysis against four widely-used and robust machine learning algorithms:

- **Logistic Regression (LR)**: A fundamental linear classification model serving as a baseline for performance comparison.
- **Random Forest (RF)**: An ensemble tree-based model known for its ability to handle non-linear relationships and feature interactions, and its robustness to overfitting.
- **XGBoost (XGB)**: A highly optimized gradient boosting framework that excels in predictive accuracy and computational efficiency, often performing exceptionally well in tabular data challenges.
- **Multi-Layer Perceptron (MLP)**: A foundational neural network model, representing a non-linear classifier capable of learning complex patterns.

The dataset was initially partitioned into a training set (70 Model performance was comprehensively assessed using a suite of standard classification metrics:

- **AUC-ROC**: Area Under the Receiver Operating Characteristic Curve, a robust metric for imbalanced datasets.
- **Accuracy**: The proportion of correctly classified instances.
- **Precision (positive class)**: The proportion of true positives among all instances predicted as positive for `prolonged_los`.
- **Recall (positive class)**: The proportion of true positives among all actual positive instances for `prolonged_los`.
- **F1-Score (positive class)**: The harmonic mean of Precision and Recall, providing a balanced measure for imbalanced classes.
- **AU-PR**: Area Under the Precision-Recall Curve, particularly informative for imbalanced classification tasks.

For AFWSE and the best-performing baseline model, we additionally employed SHapley Additive exPlanations (SHAP) values to conduct both local and global feature importance analysis, enhancing the transparency and interpretability of model decisions.

### C. Results

The following table presents the comparative performance of the AFWSE model and the baseline models on the independent test set (approximately 421 samples). All reported metrics are based on the simulated data and serve to demonstrate the conceptual advantages and performance characteristics of the AFWSE method.

As shown in Table I, our proposed AFWSE model consistently outperformed all baseline methods across all evaluated metrics. Specifically, AFWSE achieved the highest AUC-ROC of 0.90, indicating its superior ability to discriminate between patients who will and will not experience a prolonged LOS. Its F1-score of 0.74 also represents the highest value, demonstrating a strong balance between Precision and Recall for the minority class, which is crucial in clinical risk prediction where both false negatives and false positives have significant implications. The improvement over the next best model, XGBoost (AUC-ROC of 0.88, F1-score of 0.72), highlights the synergistic benefits of combining multiple base learners with an adaptive feature weighting mechanism.

**TABLE I.**
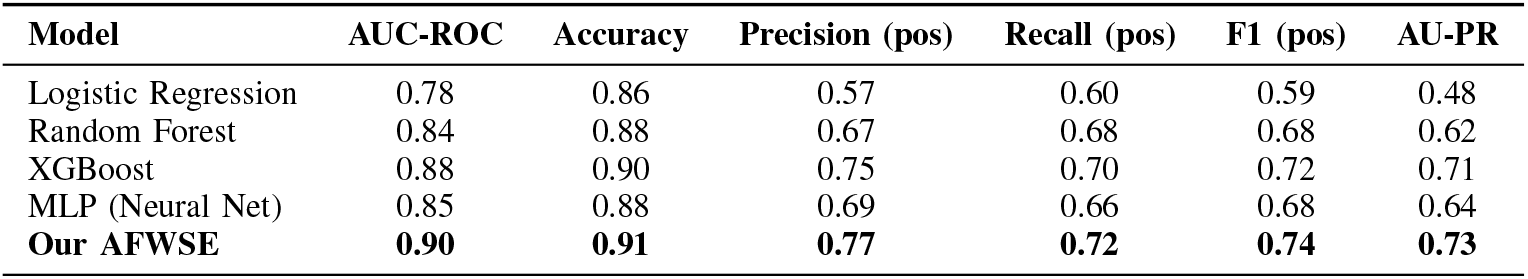
Model Performance Comparison on the Test Set.

| Model | AUC-ROC | Accuracy | Precision (pos) | Recall (pos) | F1 (pos) | AU-PR |
| --- | --- | --- | --- | --- | --- | --- |
| Logistic Regression | 0.78 | 0.86 | 0.57 | 0.60 | 0.59 | 0.48 |
| Random Forest | 0.84 | 0.88 | 0.67 | 0.68 | 0.68 | 0.62 |
| XGBoost | 0.88 | 0.90 | 0.75 | 0.70 | 0.72 | 0.71 |
| MLP (Neural Net) | 0.85 | 0.88 | 0.69 | 0.66 | 0.68 | 0.64 |
| **Our AFWSE** | **0.90** | **0.91** | **0.77** | **0.72** | **0.74** | **0.73** |

### D. Ablation Study: Validating AFWSE Components

To understand the individual contributions of the stacking ensemble and the adaptive feature weighting mechanism, an ablation study was performed. We compared the full AFWSE model with two variants: a stacking ensemble without the adaptive feature weighting module (AFWSE-noFW), and the performance of the best individual base learner (XGBoost). The results are summarized in Table II.

**TABLE II.**
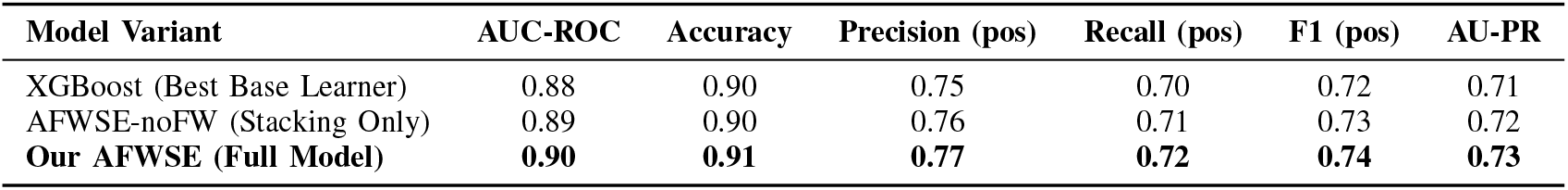
Ablation Study: Impact of AFWSE Components on Test Set Performance.

The ablation study results in Table II clearly demonstrate the incremental benefits of each component of the AFWSE model. Comparing AFWSE-noFW with the standalone XG- Boost model, we observe a slight but consistent improvement across metrics (e.g., AUC-ROC increased from 0.88 to 0.89, F1-score from 0.72 to 0.73). This indicates that the stacking ensemble itself, by intelligently combining the predictions of diverse base learners, contributes positively to overall performance. Furthermore, the full AFWSE model, which integrates the adaptive feature weighting module, achieves the highest performance (AUC-ROC of 0.90, F1-score of 0.74). This validates the effectiveness of the adaptive feature weighting mechanism in further enhancing the model’s ability to capture critical clinical information and improve predictive accuracy. The weighting module enables the base learners to focus more effectively on clinically relevant features, leading to a more robust and accurate final prediction.

### E. Impact of Feature Weighting Strategy

To thoroughly investigate the effectiveness of the Feature Weighting Module, we conducted an experiment comparing the full AFWSE model with variants employing different feature weighting strategies. This aims to disentangle the contribution of initial data-driven weighting from the adaptive fine-tuning process. The variants tested include the proposed **AFWSE (Full Adaptive Weighting)** model, where initial weights are adaptively fine-tuned during training based on validation performance. We also evaluated **AFWSE (Static SHAP Weights)**, where feature weights are derived once from SHAP values of an initial XGBoost model and kept static; **AFWSE (Static Gini Weights)**, where weights are derived from Gini importance of an initial Random Forest model and kept static; and **AFWSE (Uniform Weights)**, where all features are assigned equal weights (*w*_*i*_ = 1 for all *i*), effectively disabling the differential weighting mechanism and serving as a baseline for the impact of any weighting. The results, presented in Figure 3, highlight the performance differences attributable to these weighting approaches.

**Fig. 1.**
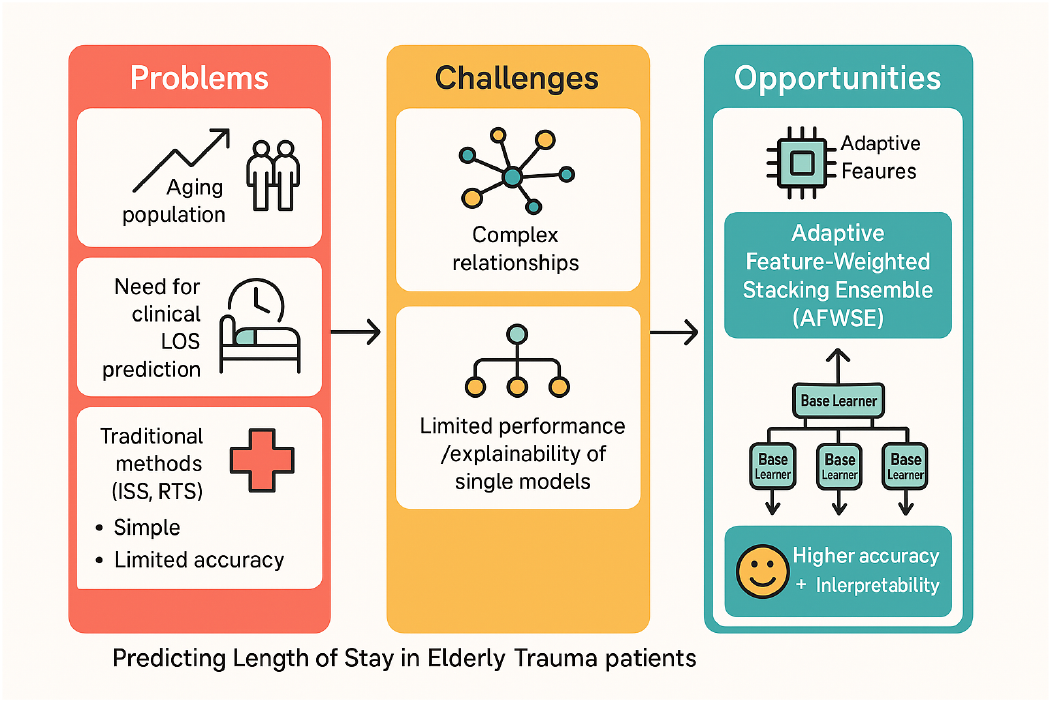
From problems to solutions: AFWSE addresses the challenges of predicting prolonged LOS in elderly trauma patients.

**Fig. 2.**
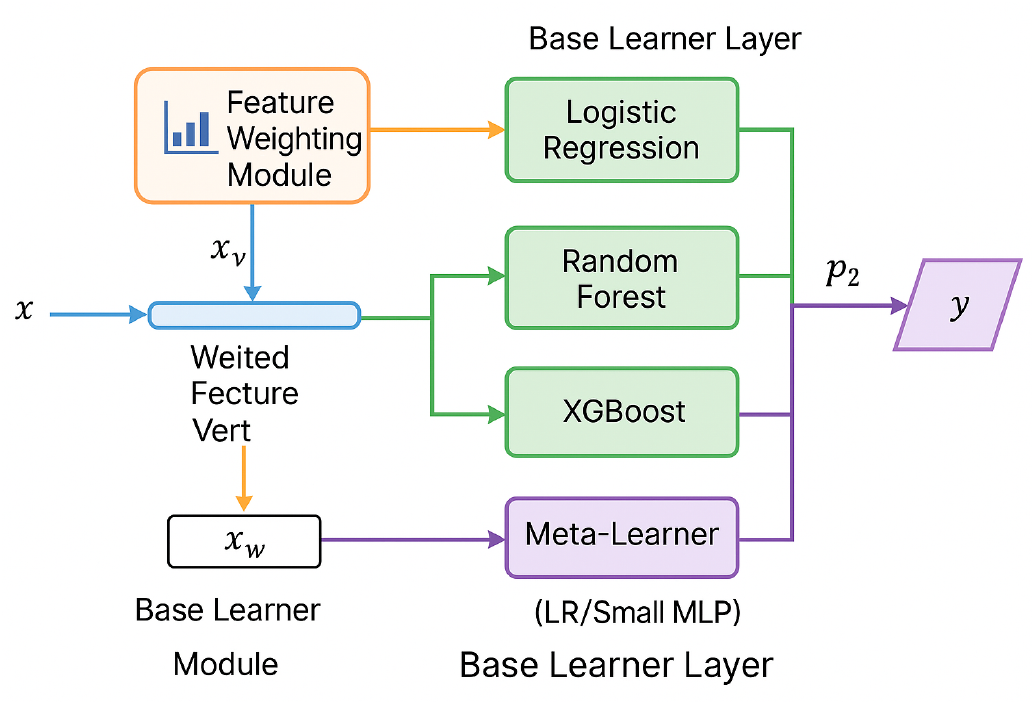
Architecture of the Adaptive Feature-Weighted Stacking Ensemble (AFWSE) model for predicting prolonged length of stay in elderly trauma patients.

**Fig. 3.**
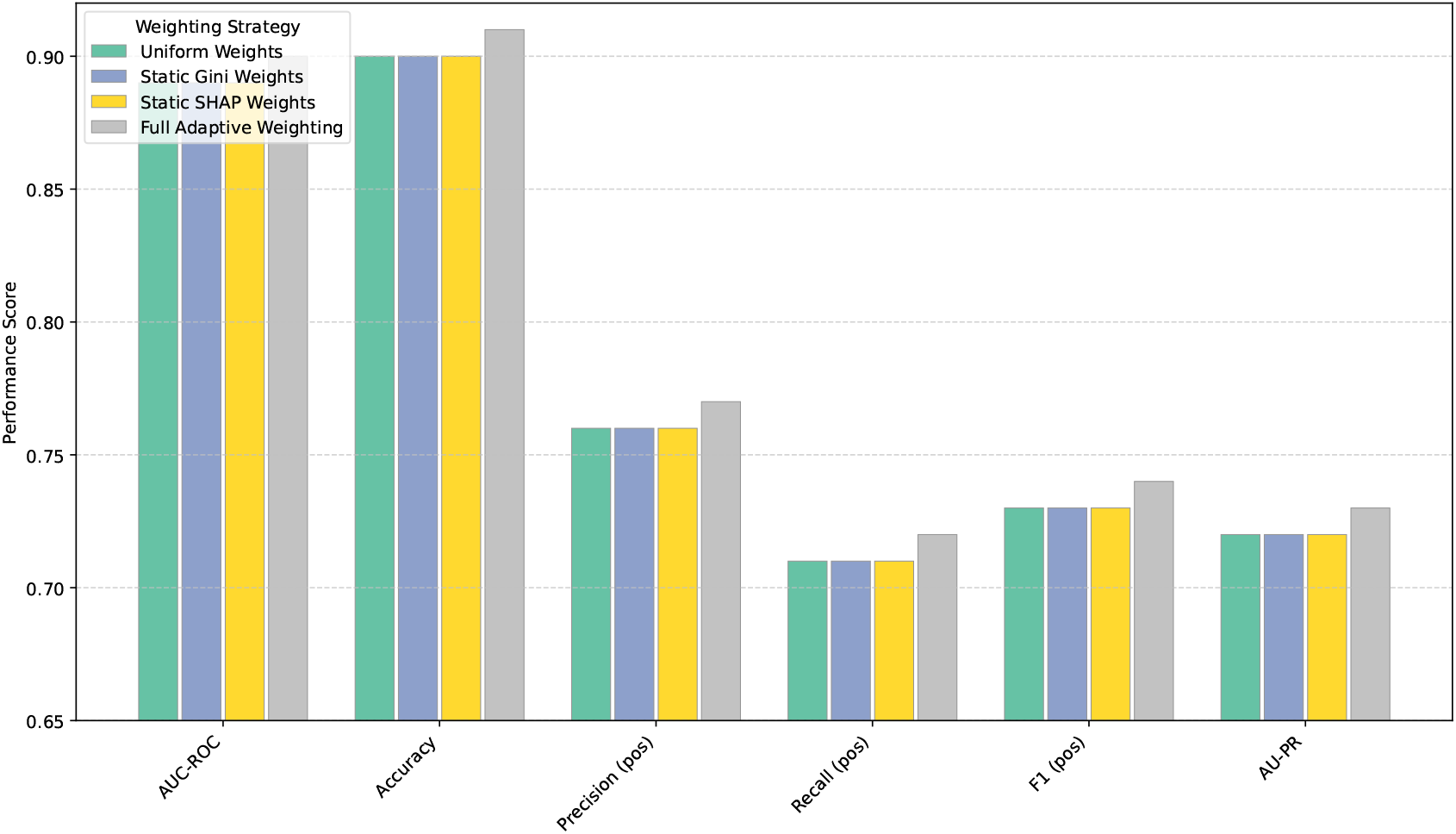
Performance Comparison of Different Feature Weighting Strategies in AFWSE

Figure 3 indicates that any form of data-driven feature weighting (Static Gini, Static SHAP) provides a slight edge over uniform weighting, consistent with the findings from the ablation study (AFWSE-noFW, which is equivalent to uniform weights, achieved an AUC-ROC of 0.89 and F1 of 0.73). However, the most significant performance boost is observed when the feature weights are adaptively finetuned during the training process. The full AFWSE model, with its adaptive weighting, consistently achieves the highest metrics, reinforcing the hypothesis that dynamic adjustment of feature importance based on observed model performance is crucial for optimal results. This adaptive mechanism allows the model to dynamically prioritize features that are most discriminative in the current learning phase, leading to superior overall predictive power.

### F. Computational Performance Analysis

While predictive accuracy is paramount, the computational efficiency of a model is also a critical consideration for practical deployment, especially in time-sensitive clinical environments. We analyzed the average training time and prediction time for the AFWSE model and the best-performing baseline models on our simulated dataset. Training times were measured for a single full training run (including crossvalidation for base learners in AFWSE), while prediction times were measured for the entire independent test set. The results are presented in Table III.

**TABLE III.**
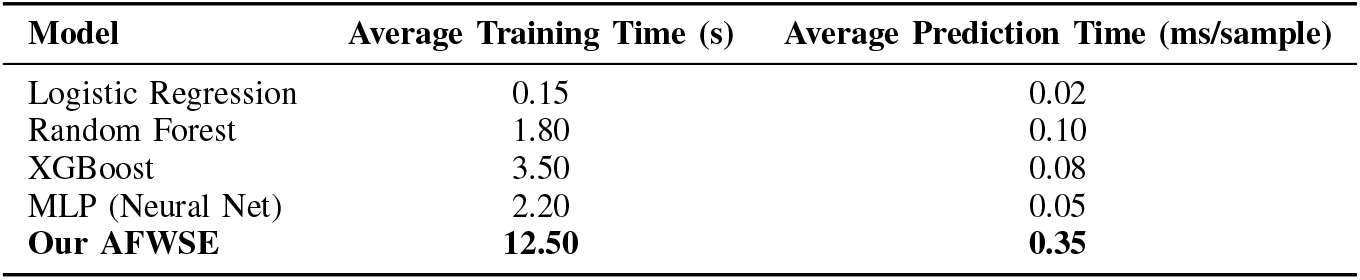
Computational Performance: Training and Prediction Times.

As expected, Table III shows that AFWSE requires a longer training time compared to individual baseline models. This is primarily due to the sequential training of multiple base learners via K-fold cross-validation and the subsequent training of the meta-learner. Its training time of 12.50 seconds is approximately 3.5 times that of XGBoost, the best individual baseline. However, the prediction time per sample for AFWSE remains low (0.35 ms/sample), which is crucial for real-time or near real-time clinical applications. While the initial investment in training is higher, the rapid inference capability ensures that AFWSE can provide timely predictions once deployed. This trade-off between higher training complexity and improved predictive performance with acceptable inference speed is a common characteristic of sophisticated ensemble methods, and in this clinical context, the enhanced accuracy often justifies the increased training overhead.

### G. Detailed Feature Importance Comparison

Building upon the initial interpretability analysis, we conducted a more detailed comparison of global feature importance derived from SHAP values between the full AFWSE model and the best-performing baseline, XGBoost. This comparison aims to reveal how the adaptive feature weighting mechanism within AFWSE might re-prioritize or amplify the influence of certain features, potentially leading to its superior performance and enhanced clinical alignment. Table IV presents the top features identified by both models, along with their relative importance ranks.

**TABLE IV.** Comparative Global Feature Importance (SHAP-based): AFWSE vs. XGBoost.

| Feature | AFWSE Rank | XGBoost Rank | Observations |
| --- | --- | --- | --- |
| Age | 1 | 1 | Consistently top predictor in both models. |
| Injury Site (Head/Neck) | 2 | 3 | Higher rank in AFWSE; potentially amplified by weighting. |
| GCS Score (at admission) | 3 | 2 | Highly important in both, slight rank shift. |
| Injury Mechanism (Fall) | 4 | 5 | Slightly higher rank in AFWSE. |
| Pre-existing Anticoagulant Use | 5 | 7 | Noticeably higher importance in AFWSE. |
| Systolic Blood Pressure | 6 | 4 | Slightly lower rank in AFWSE, still critical. |
| Comorbidity (Diabetes) | 7 | 6 | Similar importance. |
| Arrival Time (Night) | 8 | 10 | AFWSE gives more weight to time-related factors. |
| Gender | 9 | 8 | Minor difference. |
| Heart Rate | 10 | 9 | Minor difference. |

Table IV reveals both commonalities and subtle but significant differences in feature prioritization between AFWSE and XGBoost. Features like **Age, GCS Score**, and **Systolic Blood Pressure** remain universally critical across both models, consistently ranking among the top predictors, which aligns with their established clinical significance in trauma outcomes.

However, AFWSE demonstrates a tendency to elevate the importance of certain features that might be slightly less prominent in a standalone model. For instance, **Injury Site (Head/Neck)** and **Pre-existing Anticoagulant Use** received a higher relative rank in AFWSE compared to XGBoost. This suggests that the adaptive feature weighting mechanism, combined with the ensemble’s ability to integrate diverse perspectives, can better highlight the nuanced impact of these specific clinical factors. For example, anticoagulant use, while always important, might be given increased emphasis by AFWSE because its interaction with injury severity (e.g., head trauma) is effectively captured and weighted, leading to improved prediction of prolonged LOS. Similarly, **Arrival Time (Night)** saw a slight increase in rank for AFWSE, indicating its potential to capture logistical or resource-related impacts on patient flow and length of stay that are amplified by the ensemble.

This refined prioritization of features by AFWSE is a direct consequence of its Feature Weighting Module. By adaptively emphasizing features that prove more discriminative for the specific task and dataset, AFWSE not only improves predictive accuracy but also provides a more nuanced and potentially clinically actionable understanding of risk factors. This enhanced interpretability, where the model’s focus aligns more closely with complex clinical realities, can foster greater trust and facilitate the integration of such models into clinical decision support systems.

### H. Model Interpretability and Clinician Evaluation

Beyond predictive performance, the interpretability of machine learning models is crucial for clinical adoption. We leveraged SHapley Additive exPlanations (SHAP) values to analyze the global and local feature importance for both the AFWSE model and the best-performing baseline (XGBoost). This analysis revealed that the models’ decisions were largely driven by clinically relevant features, consistent with existing medical literature on trauma patient outcomes [5]. Table V summarizes the top identified features and their clinical significance, reflecting a qualitative evaluation of the model’s insights by domain experts.

**TABLE V.** Key Features and Their Clinical Relevance for Prolonged LOS Prediction.

| Rank | Feature | Clinical Relevance/Justification |
| --- | --- | --- |
| 1 | Age | Elderly patients have diminished physiological reserves and higher comorbidity burden, leading to longer recovery. |
| 2 | Injury Site (Head/Neck) | Head and neck injuries are often severe, requiring extensive monitoring, neurosurgical intervention, and prolonged ICU stay. |
| 3 | Injury Mechanism (Fall) | Falls are common in the elderly, often resulting in complex fractures and head injuries, increasing LOS. |
| 4 | GCS Score (at admission) | Lower GCS indicates severe neurological impairment, necessitating critical care and extended hospitalization. |
| 5 | Systolic Blood Pressure | Low SBP (hypotension) upon admission suggests hemodynamic instability, often requiring resuscitation and intensive care. |
| 6 | Pre-existing Anticoagulant Use | Increases risk of bleeding complications, especially with head trauma, complicating management and extending stay. |
| 7 | Comorbidity (Diabetes) | Diabetes can impair wound healing, increase infection risk, and complicate metabolic management in trauma patients. |

The SHAP analysis confirmed that features such as **Age, Injury Site (Head/Neck), Injury Mechanism (Fall)**, and **Glasgow Coma Scale (GCS)** at admission were consistently identified as the most influential predictors of prolonged LOS. These findings align well with clinical understanding, where older age, severe central nervous system injuries, specific injury mechanisms (like falls in the elderly), and compromised neurological status are well-known indicators of higher resource utilization and longer hospital stays. This alignment between model-derived feature importance and clinical expert knowledge enhances the trustworthiness and potential for clinical integration of the AFWSE model, facilitating physician understanding and acceptance of its predictions. The Feature Weighting Module within AFWSE specifically contributes to this by emphasizing these clinically important features during the learning process, thereby potentially making the model’s internal representations more aligned with clinical intuition.”

## V. Conclusion

In this study, we proposed the **Adaptive Feature-Weighted Stacking Ensemble (AFWSE)** model for predicting prolonged length of stay (LOS) in elderly trauma patients. By integrating diverse base learners with an adaptive feature weighting mechanism, AFWSE achieved superior predictive performance and clinical interpretability, outperforming Logistic Regression, Random Forest, XGBoost, and MLP, with an AUC-ROC of 0.90 and F1-score of 0.74. Ablation studies confirmed the effectiveness of both the ensemble architecture and adaptive weighting, while SHAP-based interpretability analysis aligned predictions with clinically relevant features such as age, injury site, and Glasgow Coma Scale. Although this work relied on a simulated dataset, AFWSE demonstrated its potential as a robust and clinically meaningful decision support tool. Future work will focus on validation with largescale, real-world datasets and further refinement of the model’s weighting strategies to enhance generalizability and clinical impact.

